# Structure of human lymphoid-specific helicase HELLS in its autoinhibitory state

**DOI:** 10.64898/2025.12.06.692698

**Authors:** Gundeep Kaur, Ren Ren, Jisun Lee, John R. Horton, Xing Zhang, Yang Gao, Taiping Chen, Xiaodong Cheng

## Abstract

<u>Hel</u>icase, <u>L</u>ymphoid <u>S</u>pecific (HELLS), also known as <u>L</u>ymphoid-<u>S</u>pecific <u>H</u>elicase (LSH), is a member of the SNF2 chromatin-remodeling family that regulates DNA methylation and heterochromatin organization. Unlike most chromatin remodelers, HELLS is catalytically inactive in its apo form and requires the DNA-binding protein CDCA7 for activation, though the underlying mechanism has remained unclear. Here, we combine biochemical, biophysical and cryo-electron microscopy analyses to define the structural basis of HELLS autoinhibition. HELLS alone assembles into a hexameric (trimer of dimers) architecture stabilized by interactions between its N-terminal coiled-coil (CC) domain and ATPase Lobe-1, while ATPase Lobe-2 remains flexible and disengaged. The CC domain functions both as an oligomerization scaffold and as an autoinhibitory module that restricts catalytic activity. Binding of CDCA7 and DNA promotes formation of an active HELLS-CDCA7-DNA ternary complex. CDCA7 recognizes hemimethylated CpG dinucleotides in both B-form and non-B-form DNA and stimulates HELLS ATPase activity. Together, these findings reveal the mechanism of HELLS autoinhibition and its activation by CDCA7 and DNA, providing new insight into how the HELLS-CDCA7-DNA ternary complex maintains DNA methylation and heterochromatin integrity.

**Graphical Abstract:** 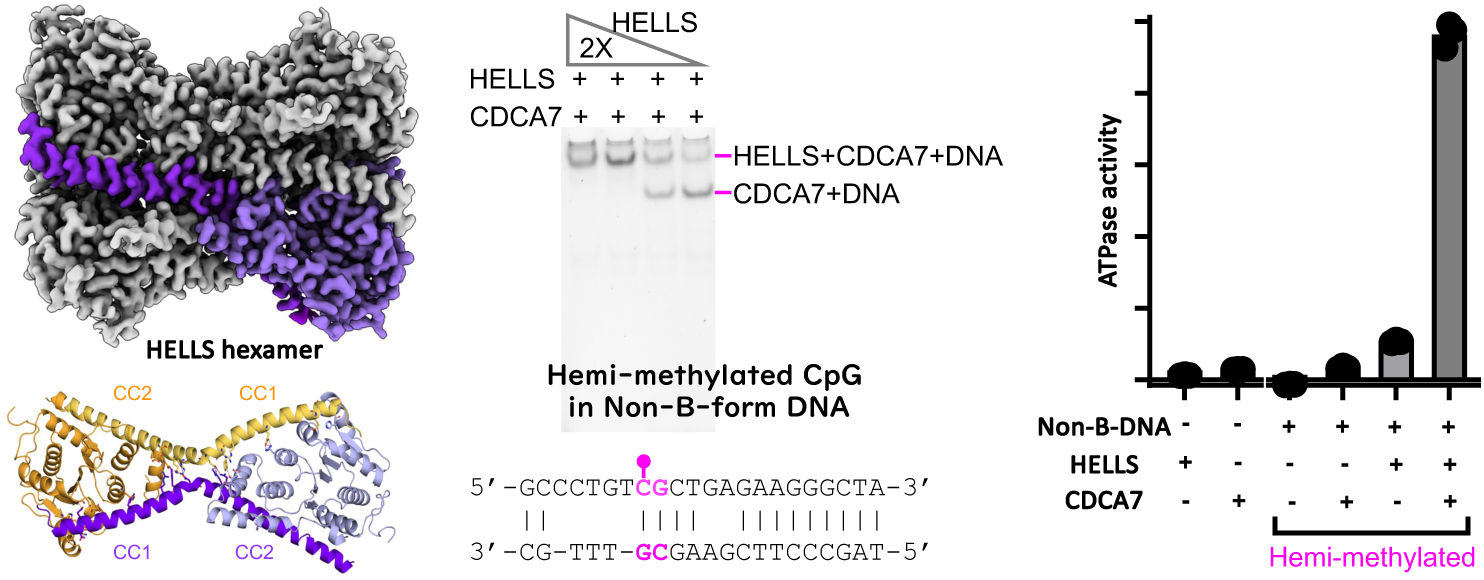

- HELLS alone assembles into an inactive hexameric trimer-of-dimers
- The N-terminal coiled-coil mediates oligomerization
- CDCA7 and DNA binding activate the HELLS-CDCA7-DNA complex
- The complex recognizes hemimethylated CpG in non-B-form DNA

## INTRODUCTION

Chromatin-remodeling complexes modulate chromatin organization and are essential for DNA-based processes such as recombination, repair, replication, and transcription (1–3). These complexes contain ATP-dependent helicases that use the energy of ATP hydrolysis to reposition, reorganize, evict, or exchange nucleosomes (4).

<u>L</u>ymphoid-<u>s</u>pecific <u>h</u>elicase (Lsh) was first identified in murine thymic precursor cells (5), and its human homolog, HELLS (<u>Hel</u>icase, <u>L</u>ymphoid <u>S</u>pecific) - also known as SMARCA6 (<u>S</u>WI/SNF2-related, <u>M</u>atrix-associated, <u>A</u>ctin-dependent <u>R</u>egulator of <u>C</u>hromatin, subfamily <u>A</u>, member <u>6</u>) or PASG (<u>P</u>roliferation-<u>A</u>ssociated <u>S</u>NF2-like <u>G</u>ene) - was cloned from an acute megakaryoblastic leukemia cell line (6). Together with the plant homolog DDM1 (<u>D</u>ecrease in <u>D</u>NA <u>M</u>ethylation <u>1</u>) (7), these proteins form a distinct subfamily within the SNF2 family of chromatin remodelers. They participate in diverse biological processes, including development, gene expression, DNA repair and tumorigenesis (8–13). Notably, HELLS has been proposed as a potential biomarker for cancer diagnosis and prognosis (14,15).

Lsh is enriched in pericentromeric heterochromatin (16), a highly compact and transcriptionally repressed region marked by DNA methylation and histone H3 lysine 9 trimethylation. Loss of *Lsh* in mice causes a 50-70% reduction in global DNA methylation, particularly at repetitive elements (17–19), a phenotype mirrored in *DDM1*-deficient *Arabidopsis* mutant (20). Thus, maintenance of DNA methylation in heterochromatin is a conserved function of this remodeler family.

*HELLS* (21) is one of five genes - along with *DNMT3B* (22–24)*, ZBTB24* (25–28)*, CDCA7* (21) and *UHRF1* (29) - mutated in Immunodeficiency, Centromeric instability, and Facial anomalies (ICF) syndrome (30–32), a rare disorder characterized by DNA hypomethylation of centromeric repeats (33). Recent studies show that ZBTB24, CDCA7 and HELLS form a regulatory pathway directing DNA methylation to specific genomic regions (34,35). ZBTB24, a C2H2 zinc-finger transcription factor, activates *CDCA7* transcription (36–38). CDCA7, a zinc-containing protein that binds hemi-methylated CpG dinucleotides (39,40), preferentially in a nonB DNA structure (41), recruits HELLS to heterochromatin to facilitate DNA methylation (34,42,43).

Unlike the plant homolog DDM1 (44), purified Lsh and HELLS lack detectable ATPase and nucleosome-sliding activity (11,43,45). HELLS contains an N-terminal coiled-coil (CC) domain and two RecA-like ATPase lobes (Fig. 1A). The CC domain acts as an autoinhibitory element, preventing ATPase activity and nucleosome remodeling in the apo form (39). Autoinhibition can be relieved by CC deletion or by binding of CDCA7 and DNA (39). However, the molecular mechanism underlying HELLS autoinhibition and activation remains unclear. Here, we report the 2.86-Å cryo-electron microscopy (cryo-EM) structure of HELLS in its autoinhibited state. In the apo form, HELLS assembles as a hexamer, with the CC domain mediating oligomerization. Upon binding CDCA7 and DNA, HELLS forms an active HELLS-CDCA7-DNA ternary complex that exhibits ATPase activity.

**Figure 1.**
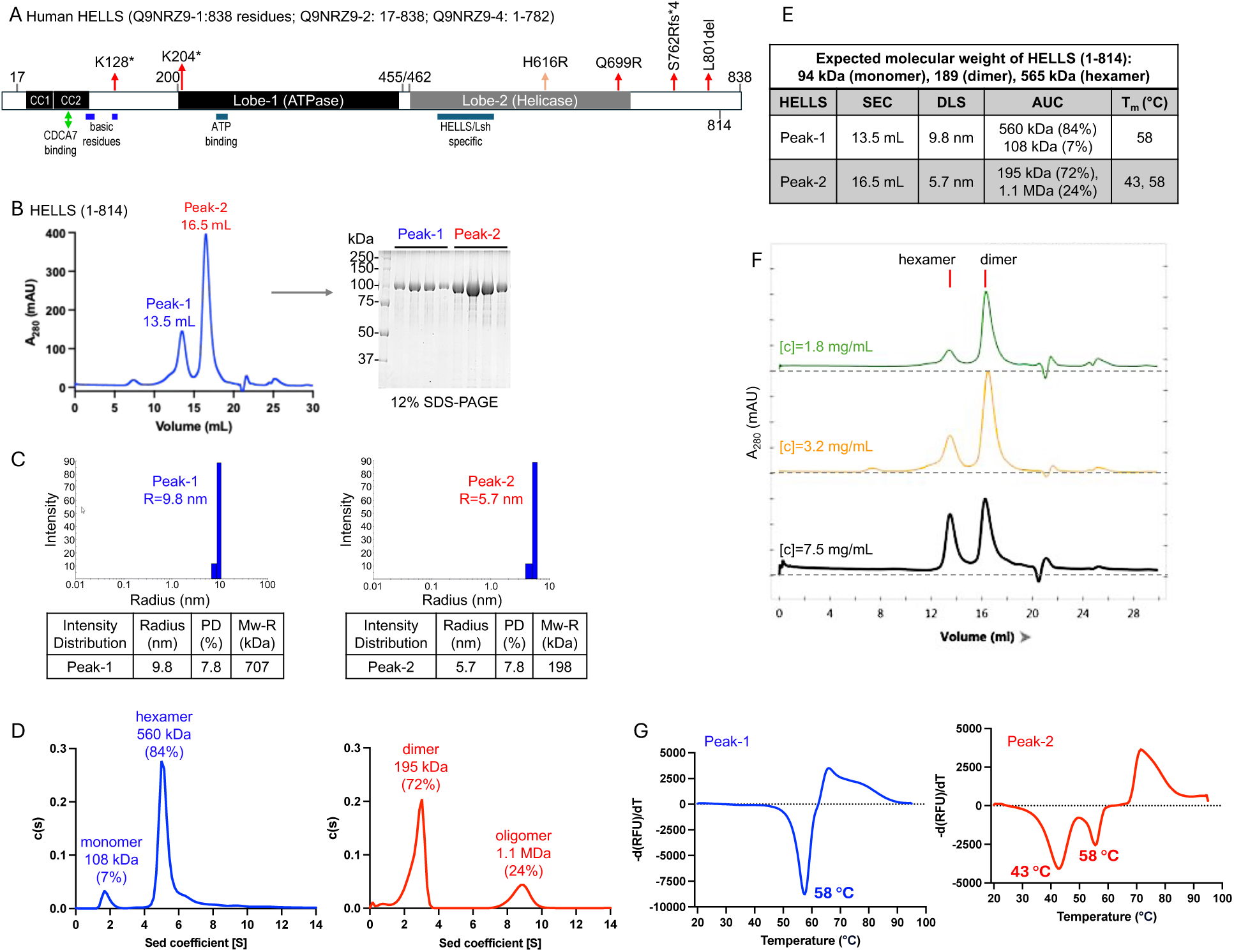
Purification and biophysical characterization of recombinant HELLS. (**A**) Domain organization of human HELLS showing the N-terminal coiled-coil (CC) domain and ATPase Lobe-1 and Lobe-2. The lengths of the three isoforms and their corresponding UniProt accession numbers are provided. Known ICF-associated mutations are marked above the sequence. (**B**) SEC profile on a Superose6 increase 10/300GL column showing two major peaks of HELLS (1-814) with distinct elution volumes. SDS-PAGE analysis of fractions from both peaks is shown to the right. (**C**) DLS analysis of individual peak fractions, showing their hydrodynamic radii. (**D**) SVAUC sedimentation profiles indicating the molecular masses of the major and minor species. (**E**) Summary of biophysical parameters for the two peaks, including SEC elution volumes, DLSderived hydrodynamic radii, AUC-derived molecular masses, and melting temperatures. (**F**) SEC profile demonstrating concentration-dependent hexamer formation. (**G**) Thermal melting curves showing the stability and melting temperatures of each peak.

## MATERIAL AND METHODS

### Expression and purification of recombinant HELLS

We generated several human HELLS constructs and used three in this study, full-length (residues 1-838; pXC2259), a C-terminal truncation (residues 1-814; pXC2274), and a Lobe-2 deletion (residues 1-460; pXC2266). All were cloned into a modified pET28b vector (pETHisSumo) with an N-terminal 6×His-SUMO tag and expressed in *Escherichia coli* BL21-CodonPlus (DE3)-RIL. Cultures were grown in LB medium initially at 37°C until OD_600_ of 0.4-0.5, and cooled to 16°C. Protein expression was induced, until OD_600_ reached ∼0.8, with 0.2 mM isopropyl β-D-1thiogalactopyranoside (IPTG) for 20 h at constant temperature of 16°C.

Cells were harvested and lysed at 4°C in 20 mM Tris-HCl (pH 7.5), 500 mM NaCl, 5% glycerol, and 0.5 mM tris(2-carboxyethyl)phosphine (TCEP), followed by sonication. After centrifugation (47,000×g for 50 min), the cell lysate was loaded onto a 5 mL HisTrap HP column (Cytiva), washed with 20 mM imidazole, and eluted using a 20–500 mM imidazole gradient. Pooled fractions were digested with home-made ULP1 protease to remove the His-SUMO tag.

The cleaved sample was applied to a HiTrap SP HP cation exchange column (Cytiva) and eluted with a 0.1–1 M NaCl gradient in 20 mM Tris-HCl (pH 7.0), 5% glycerol, and 0.5 mM TCEP. Peak fractions were pooled and reapplied to the HisTrap HP column to remove residual contaminants, then concentrated and further purified by size-exclusion chromatography (SEC) on a Superdex 200 16/60 column (Cytiva) equilibrated in 20 mM Tris-HCl (pH 7.5), 150 mM NaCl, and 0.5 mM TCEP. Two major peaks were observed, visualized by SDS-PAGE and Coomassie staining. Fractions were collected separately, concentrated, flash-frozen in liquid nitrogen, and stored at –80°C for later use.

In addition, purified and concentrated HELLS proteins (residues 1-838, 1-814, and 1-460), along with a set of five molecular-weight calibration standards (Bio-Rad 1511901), were analyzed by analytic SEC column using a Superose 6 Increase 10/300 GL (Cytiva) in 20 mM Tris-HCl (pH 7.5), 150 mM NaCl, and 0.5 mM TCEP at 4°C. Samples were injected at the flow rate of 0.5 mL/min. For the pXC2274 construct (residues 1-814), samples were run at multiple protein concentrations to determine its elution profile and oligomeric states.

### Expression and purification of CDCA7

We generated a full-length human CDCA7 construct (residues 1-371; pXC2402) and cloned it into the baculovirus transfer vector pVL1393-6xMyc (Addgene 128035) with an N-terminal HisMBP tag and a C-terminal 6xMyc tag. The 6xMyc tag allowed us to verify the integrity of the Cterminus. Recombinant baculovirus was produced by co-transfecting Sf9 cells with a linearized BestBac 2.0 Δv-cath/chiA genome (Expression Systems) using Cellfectin II (Thermo Fisher). For protein expression, Sf9 cells were cultured in suspension in Sf-900 II SFM medium supplemented with antibiotic-antimycotic (Gibco) and 0.5% heat-inactivated fetal bovine serum. At a density of 1.3 x 10^6^ cells/mL, cultures were infected with 3% (v/v) baculovirus and incubated at 27℃ for 72 h. Cells were harvested and lysed by sonication at 4 ℃ in 20 mM Tris-HCl (pH 8.0), 500 mM NaCl, 5% glycerol, 0.5 mM TCEP, 100 µM phenylmethylsulfonyl fluoride (PMSF) and an EDTAfree protease inhibitors (Pierce catalog #A32965). The clarified lysate, after centrifugation at 44,000 × g for 30 min at 4℃, was filtered (0.45 μm PVDF membrane) and loaded onto a 1-mL HisTrap HP column (Cytiva), washed with 30 mM imidazole, and eluted by stepwise imidazole gradient. Pooled fractions were further purified on a 1-mL HiTrap™ ANX Sepharose FF column (Cytiva) using a 0.2-1 M NaCl gradient in 20 mM Tris-HCl (pH 8.0), 5% glycerol, and 0.5 mM TCEP. Final fractions were concentrated, flash-frozen in liquid nitrogen, and stored at -80℃ for later use.

To assemble the CDCA7-HELLS complex, purified CDCA7 and HELLS were mixed at a molar ratio of 2:1 in 20 mM Tris-HCl (pH 8.0), 250 mM NaCl, 5% Glycerol, and 0.5 mM TCEP, and incubated for 1 h at 4℃. The mixture was then loaded onto a Superdex 200 10/300 GL column (Cytiva) equilibrated in the same buffer and run at 0.3 mL/min flow rate at 4℃.

### Dynamic light scattering (DLS)

Fractions of purified HELLS proteins (residues 1-814) at hexamer and dimer peaks were adjusted to ∼0.3 mg/ml, and 20 μL of each sample was transferred into a 384-well plate. The plate was centrifuged at 3000xg for 5 min at 4°C and then loaded into a DynaPro Plate Reader-II (Wyatt Technology) for conducting dynamic light scattering analysis. Data were collected for 10s per well, and results were analyzed and plotted using DYNAMICS 7.1.9 software (https://www.wyatt.com/products/software/dynamics.html).

### Sedimentation velocity analytical ultracentrifugation (SV-AUC)

Purified HELLS proteins - dimer and hexamer fractions of the residues 1-814 construct (0.3 mg/mL each) and the residues 1-460 construct (0.15 mg/mL) - were dialyzed overnight at 4°C in 20 mM Tris (pH 7.5), 150 mM NaCl, 5% glycerol, and 0.5 mM TCEP. A Beckman-Coulter XL-A ultracentrifuge equipped with An-60 Ti four-hole rotor was pre-cooled and equilibrated at 4°C in the vacuum chamber for ∼2 h before use. Sedimentation was monitored by absorbance at 280 nm at 1-min intervals during centrifugation at 40,000 rpm at 4°C. The partial specific volume, solvent density, and viscosity were calculated using SEDNTERP (46), and the data were analyzed with SEDFIT (47) to obtain c(s) distribution. Final plots were prepared using GraphPad Prism 9.0 (https://www.graphpad.com/features).

### Differential scanning fluorimetry (thermal shift assay)

Thermal shift assays were performed using SYPRO Orange dye (Invitrogen S6651) on a CFX Opus 96 Real-Time PCR System (#12011319). Purified HELLS proteins (dimer and hexamer of residues 1-814 at 2 μM) were mixed with SYPRO Orange dye (5x final concentration), 1 mM MgCl_2_ and 20 μM ATPγS in a 20 µL reaction. Samples were heated from 20°C to 95°C at a rate of 0.5°C per min. Melting curves and data analysis were generated using CFX Maestro software (https://www.bio-rad.com/en-us/product/cfx-maestro-software-for-cfx-real-time-pcr-instruments), and the final plots were prepared in GraphPad Prism 9.0 (https://www.graphpad.com/features).

### Electrophoretic mobility shift assay (EMSA)

Purified CDCA7, HELLS, or the CDCA7-HELLS mixture was incubated with DNA (50 nM) in 20 mM Tris-HCl pH8, 150 mM NaCl, 5% glycerol, 0.5 mM TCEP for 1 h on ice. Samples were resolved on an 8% native polyacrylamide gel at 150 V for 45 min in ice-cold 0.5× TBE. Gels were stained with SYTOX Green (catalog no. S7020) at a 1:20,000 dilution in water at room temperature for 10 min and imaged on a Typhoon 9410 variable mode imager (GE Healthcare) on the Cy2 channel.

### ATPase activity

ATPase activity was measured using the ADP-Glo Kinase Assay Kit (Promega catalog #V9101). Reactions (20 μL) contained 20 mM Tris-HCl (pH 8.0), 120 mM NaCl, 4 mM MgCl_2_, 0.4 mM ATP, 1 mM DTT and 0.5 mg/ml bovine serum albumin and were incubated at 37°C for 1.5 h followed by 30 min at room temperature. For analysis, 5 μL of each reaction was transferred into a 384-well plate (Corning #4513) in triplicate. ADP-Glo Reagent (5 μL) was added and incubated for 40 min at room temperature, followed by 10 μL of Kinase Detection Reagent and 1-h incubation. Luminescence was recorded using a Synergy Neo2 Multi-mode Reader (BioTek) and converted to ADP concentration using a linear standard curve.

### Cryo-EM sample preparation and data collection

Purified HELLS (residues 1-814) dimer and hexamer samples (2 μM) were mixed with 1 mM MgCl_2_ and 20 μM ATPγS and incubated on ice for ∼30 min. Quantifoil R1.2/1.3 200 mesh Cu grids were glow-discharged at 20 mA for 30s using the ‘air negative, positive surface’ setting on the GloQube Plus. Each sample (3.5 μL) was applied to freshly glow-discharged grids in a Vitrobot Mark IV (Thermo Fisher Scientific) at 4°C and 100% humidity (blot force -3; blot time 3-6s), then plunge-frozen in liquid ethane. Grids were screened, and an initial data set was collected (for grid with blot time 5s) using a Titan Krios G3 microscope (Thermo Fisher Scientific) operated at 300 kV at UTHealth Cryo-Electron Microscopy Core Facility, Houston, TX.

The new grids were prepared for the best-performing condition (5s blot), and highresolution data were collected at the National Center for CryoEM Access and Training (NCCAT) and the Simons Electron Microscopy Center (New York Structural Biology Center). Data were acquired on an EF-Krios microscope operated at 300 kV with a Gatan K3 detector. Movies were recorded in super-resolution mode at 105k magnification (0.4135 Å pixel size; 0.827 Å after 2x binning) using Leginon (48). Data were collected at a dose rate of 29.26 e^-^/Å^2^/s with a 2s exposure (total dose 58.51 e^-^/Å^2^), split into 50 frames (0.04s/frame). A total of 12,544 micrographs were collected over a nominal defocus range of 0.7 – 2.0 μm.

### Cryo-EM data processing

The cryoSPRAC software v4.5.3 (49) was used for cryo-EM structure determination and nonuniform refinement, and the final reconstruction was sharpened with DeepEMhancer (50) and processed on the COSMIC2 platform (51). Two cryo-EM reconstructions were generated. The first, reconstructed with D3 symmetry, includes the N-terminal CC domain and Lobe-1 and reached a resolution of 2.86 Å (DPB 9Z05) with symmetry-expansion implemented in cryoSPARC (Supplementary Fig. S1). The second, a C1 reconstruction without imposed symmetry, captures the dynamic Lobe-2 and missing linkers (Supplementary Fig. S2-S4). Details of cryo-EM data collection and validation statistics are provided in Supplementary Table S1. Local resolutions, Fourier shell correlation, and angular distribution analyses for all cryo-EM reconstruction are provided in Supplementary Fig. S5.

### Model building, refinement, validation and structural visualization

Six copies of the N-terminal CC domain and the AlphaFold predicted model of Lobe-1 (52) were modeled and rigid-body fitted into the D3-symmetric cryo-EM map using COOT (53) and ChimeraX (54). Models were refined by real-space refinement in Phenix (55), and geometry was validated with MolProbity (56). Interface surface areas were calculated using the PISA (Proteins, Interfaces, Structures and Assemblies) server (57). Structural comparisons, superpositions, and figure preparation were performed in ChimeraX v1.9 and PyMol (Schrodinger).

## RESULTS

### Recombinant HELLS forms dimers and hexamers in solution

To characterize human HELLS, we produced two recombinant proteins in *E. coli*, full-length construct (residues 1-838) and a C-terminal truncated variant (residues 1-814). Both behaved similarly in biophysical assays, although the shorter construct showed greater solubility and more consistent purification. Size exclusion chromatography (SEC) revealed two major species for each protein, with peak-1 eluting at 13.5 mL, and peak-2 at 16.5 mL (Fig. 1B and Fig. S6A). Dynamic Light Scattering (DLS) indicated hydrodynamic radii of 9.8 nm (peak-1) and 5.7 nm (peak-2) (Fig. 1C). Sedimentation velocity-analytical ultracentrifugation (SV-AUC) estimated molecular masses of ∼560 kDa for peak-1 and 195 Da for peak-2, corresponding to predominant hexameric (84%) and dimeric (72%) species, respectively (Fig. 1D). Minor species of other molecular weights were also detected.

These results demonstrate that recombinant HELLS exists in at least two oligomeric states: a larger, earlier-eluting hexamer and a smaller, later-eluting dimer (Fig. 1E). The oligomerization was concentration-dependent, with higher protein concentrations favoring hexamer formation, as reflected by increased intensity of peak-1 (Fig. 1F). Differential scanning fluorimetry (DSF) showed that the hexameric form was more thermostable, exhibiting a melting temperature (T_m_) of 58°C, whereas the dimeric form displayed two transitions at 43°C and 58°C (Fig. 1G). The distinct T_m_ values indicate that hexamerization stabilizes HELLS, likely through increased intermolecular contacts.

### HELLS in hexameric form with D3 symmetry

To elucidate the molecular and structural basis of HELLS autoinhibition, we analyzed both hexameric and dimeric forms of the shortened HELLS construct (residues 1-814; Fig. S6B) by cryo-EM. Of the two assemblies, only the hexameric form yielded an interpretable electron density map (Fig. 2A). Below, we first describe the D3-symmetric reconstructed structure (PDB 9Z05) at the resolution of 2.86 Å (Fig. 2B). The resulting compact structure contained six protomers (chains A-F). The cryo-EM density allowed us to model residues 24-97 (corresponding to CC domain) and residues 208-457 (corresponding to ATPase Lobe-1) (Fig. 2C-2D). The first 23 residues at the N-terminus, the linker region (residues 98-207) between CC domain and Lobe-1, and the entire Lobe-2 were not resolved, likely due to conformational flexibility (see below).

**Figure 2.**
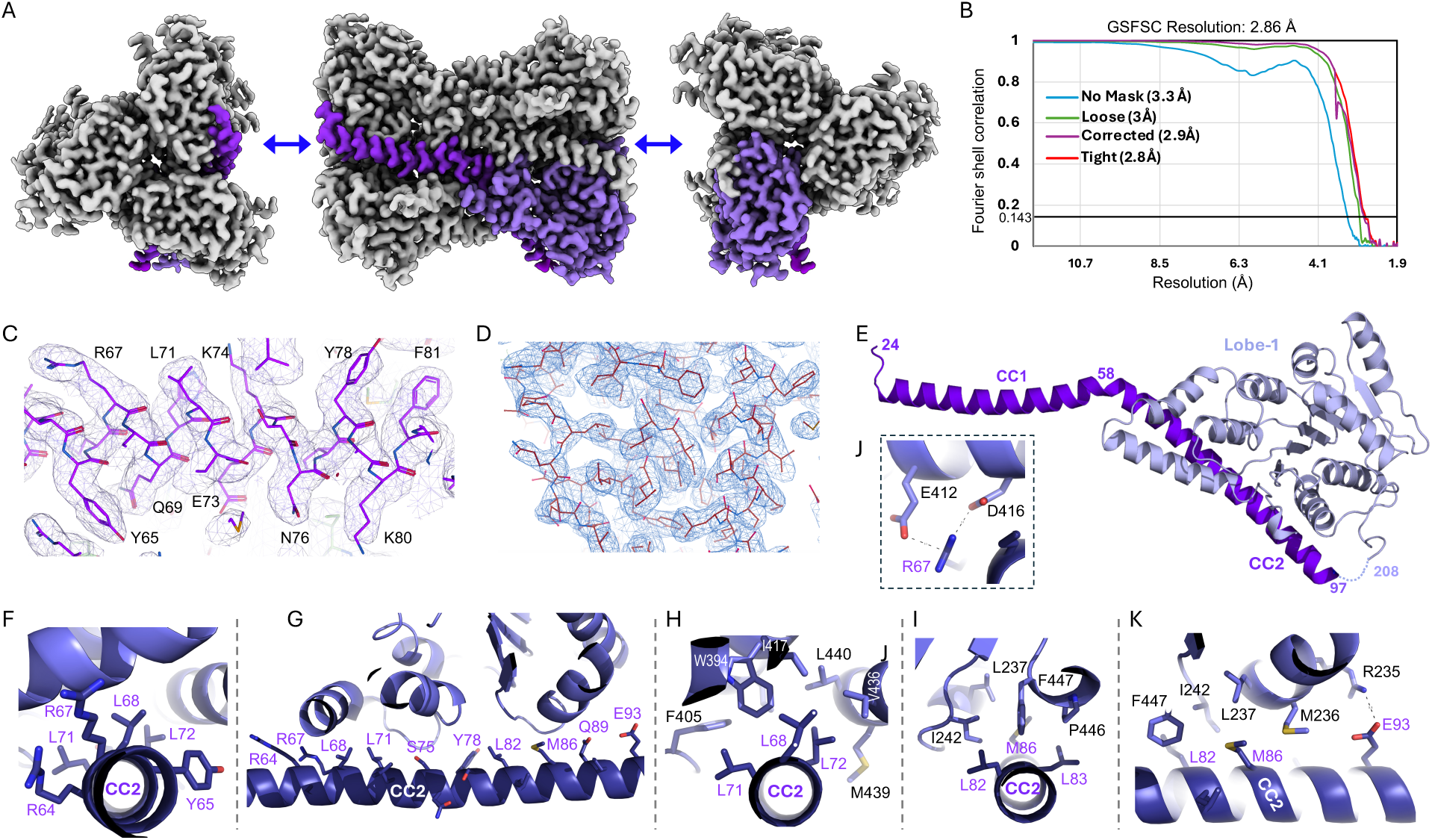
Overall architecture of HELLS hexamer and intra-molecular interactions within a HELLS protomer. **(A)** Three views of the cryo-EM density map of the D3-symmetric hexamer. One protomer is highlighted in dark blue. **(B)** Fourier shell correlation (FSC) curve between the two independently refined half-maps, indicating an overall resolution of 2.86 Å at the 0.143 cutoff. (**C-D**) Representative density maps showing well-resolved features for the CC2 helix (panel C) and Lobe-1 (panel D). (**E**) Ribbon representation of a single protomer illustrating the CC1 and CC2 helices and the seven-stranded Lobe-1. (**F**-**G**) Intra-molecular interactions mediated by CC2 residues located on one face of the helix that projects toward Lobe-1. (**H-I**) Examples of hydrophobic interactions within the interface. (**J**) Charge-charge interactions between R67 of CC2 and E412 and D416 of Lobe-1. (**K**) Coexistence of hydrophobic residues (L82 and M86) and charged residue (E93) on the same interaction face of CC2 helix.

Within each protomer, the CC domain adopts an all α-helical conformation comprising two long helices (CC1: residues 24-57 and CC2: residues 59-97) – connected by a ∼43° bend at residue 58 (Fig. 2E). ATPase Lobe-1 exhibited a mixed α/β fold characteristic of SNF2-family remodelers, consisting of a central seven-stranded parallel β-sheet flanked by multiple α-helices (58). One face of the CC2 helix makes extensive intra-molecule contacts with Lobe-1, burying an interface buried area of ∼1200 Å^2^ (Fig. 2F-2G). Two clusters of aliphatic residues, L68, L71, L72, and L82, L83, M86, located along the middle of the CC2 helix pack tightly against hydrophobic residues of Lobe1 to form the core of this interface (Fig. 2H and 2I). Charged residues at both ends of the CC2 helix contribute to electrostatic interactions, including R67 of CC2 bridging between E412 and D416 of Lobe-1 (Fig. 2J), and E93 of CC2 interacting with R235 of Lobe-1 (Fig. 2K).

### Dimer interfaces

Each protomer forms two distinct dimer interfaces within the hexamer: an A-B dimer and a B-C dimer (Fig. 3A). The A-B interface buries ∼2000 Å^2^ (Fig. 3B), whereas the B-C interface is smaller, burying ∼1400 Å^2^ (Fig. 3H). In the **A-B** dimer, Lobe-1 of each molecule is flanked by the intermolecular CC1 and intra-molecular CC2 helices (as described above) (Fig. 3B). The intermolecular interface is primarily formed by residues on one face of the CC1 helix that project toward the adjacent protomer (Fig. 3C), involving both hydrophobic and electrostatic interactions.

**Figure 3.**
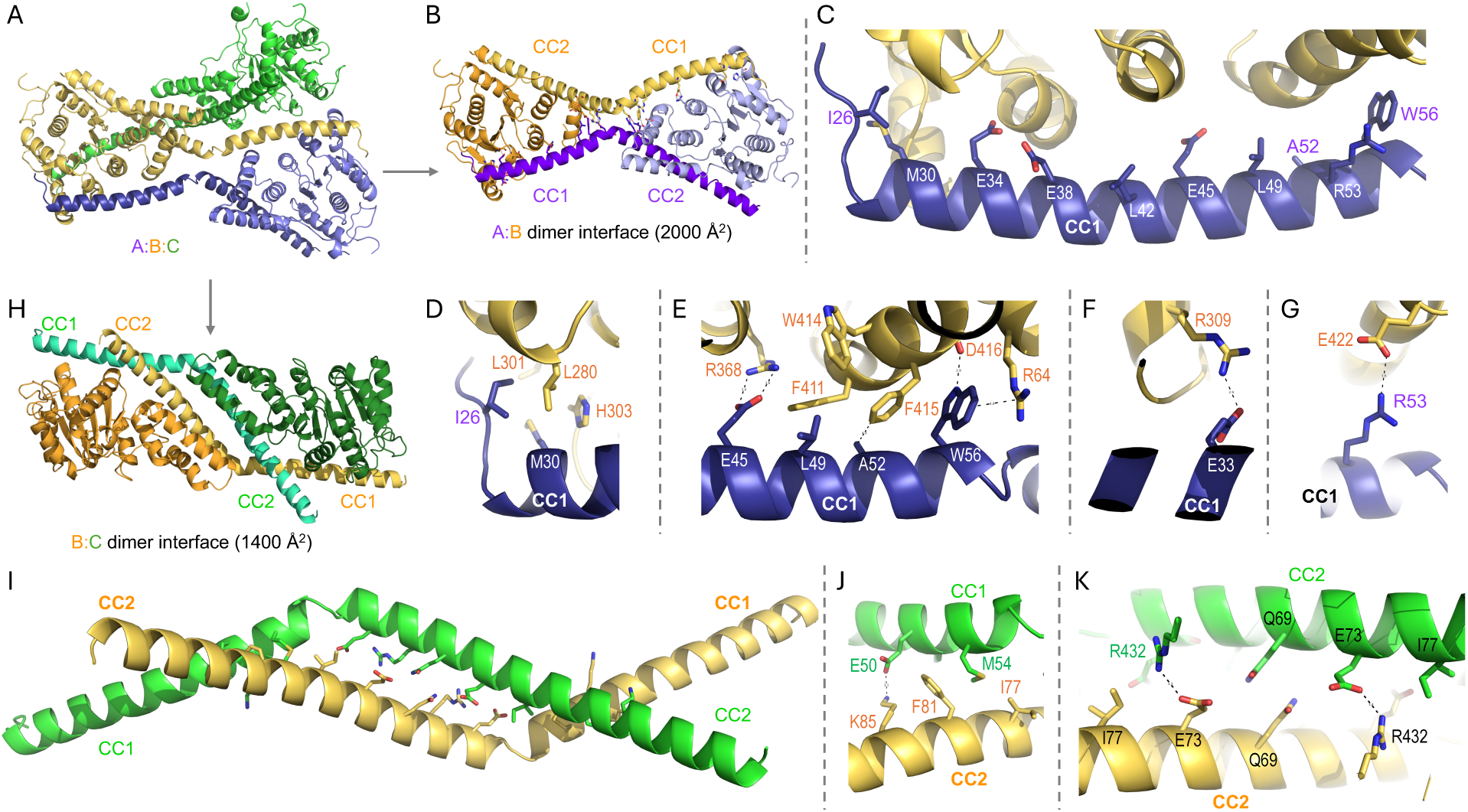
**Two distinct intermolecular dimer interfaces in the HELLS hexamer**. (**A**) Ribbons representation showing that protomer B (in yellow) forms two distinct dimer interfaces with A (blue) or C (green). (**B**) In the A-B dimer, Lobe-1 of each protomer is flanked by the intermolecular CC1 and intra-molecular CC2 helices. (**C**) Inter-molecular interactions in the A-B dimer are mediated by CC1 residues located on one face of the helix that projects toward the neighboring Lobe-1. (**D**) Hydrophobic interactions mediated by I26 and M30 of CC1. (**E**) Coexistence of charged residue E45 and hydrophobic residues L49, A52 and W56 on the same interaction face of the CC1 helix. (**F-G**) Charge-charge interactions between E33 of CC1 and R309 of Lobe-1 (panel F) and between R53 of CC1 and E422 of Lobe-1 (panel G). (**H**) In the B-C dimer, the interface is formed by the CC domains of both protomers. (**I**) CC1 and CC2 helices from the two protomers run antiparallel to form a classic coiled-coil arrangement. (**J**) CC1-CC2 interactions, including a charge-charge interaction between E50 (CC1) and K85 (CC2) and hydrophobic packing involving M54 (CC1) and I77 and F81 (CC2). (**K**) CC2-CC2 interactions, including interdigitating contacts between Q89 and E73 of CC2 from both protomers.

Hydrophobic contacts include N-terminal aliphatic residues I26 and M30 of CC1 packing against L288 and L301 of the neighboring protomer (Fig. 3D), as well as a second hydrophobic cluster formed by L49, A52, and W56 in the middle of CC1 (Fig. 3E). Three Glu-Arg salt bridges further stabilize the interface: E33 of CC1 interacts with R309 of Lobe-1 (Fig. 3F), E45 of CC1 with R368 (Fig. 3E), and R53 of CC1 with E422 (Fig. 3G). Notably, the aromatic indole ring of W56 of CC1 engages in π–π and cation–π interactions, stacking against F415 and the guanidinium group of R64 (Fig. 3E). The indole nitrogen of W56 also forms a hydrogen bond with the side-chain carboxyl oxygen of D416 (Fig. 3E).

In the **B-C** dimer, the interface is formed by the CC domains of both protomers, with their helices running antiparallel to generate a classic coiled-coil arrangement (Fig. 3I). Key interactions include an electrostatic contact between E50 of CC1 and K85 of CC2, and hydrophobic packing in which M50 of CC1 inserts between I77 and F81 of CC2 (Fig. 3J). Additionally, Q69 and E73 of CC2 from two protomers interdigitate their side chains, with E73 forming an additional stabilizing interaction with R432 of Lobe-1 (Fig. 3K).

Together, the two dimer interfaces employ both CC1 and CC2 helices to engage Lobe-1 (through intra- or inter-molecule interactions) and to form coiled-coil contacts with neighboring protomers. Importantly, residues mediating Lobe-1 interactions and those mediating coiled-coil packing occupy distinct faces of each helix. Three such dimer pairs assemble cooperatively to generate the complete hexamer.

### Tripartite interface between the trimer of dimers

We next examined the tripartite interface formed by three protomers at the center of the hexamer (Fig. 4A). This interface is dominated by interactions mediated by Lobe-1, with three Lobe-1 regions positioned at each end of the hexamer. Each protomer contributes five key residues – R338, N341, Q344, H345, and Y347 – to the interface (Fig. 4B). At the core of the assembly, Q344 residues from all three protomers form a cyclic network of hydrogen bonds that ties the trimer together (Fig. 4B). The side chain amide group of each Q344 donates a hydrogen to one neighboring protomer and the carbonyl oxygen atom accepts a hydrogen from the other, completing a three-way hydrogen-bonding ring. Electrostatic surface analysis reveals that positive electrostatic potential is concentrated at both ends of the hexamer, primarily contributed by Lobe1 residues K311, R314, K318, R319, and K210 (Fig. 4C).

**Figure 4.**
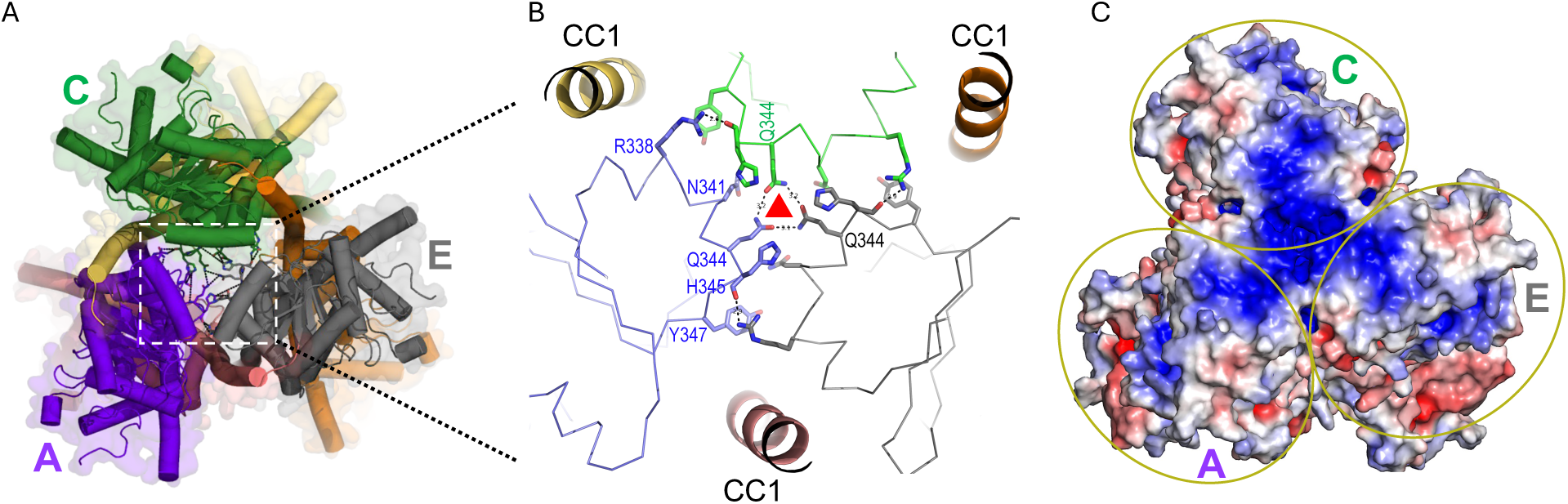
Interactions stabilizing the tripartite interface in HELLS hexamer. **(A)** HELLS hexamer viewed from opposite ends, showing the three Lobe-1 regions (protomers A, C and E) positioned at each end of the assembly. (**B**) Close-up view of the tripartite interface, highlighting a network of interactions contributed by five residues of each protomer. The central Q344 residues form a Q344 triad that links the three protomers. The CC1 helices from the remaining three protomers are shown as ribbons. (**C**) Electrostatic surface potential of the HELLS hexamer, illustrating the positively charged patch formed at the ends of the hexamer.

Since the D3 symmetric hexamer reconstruction lacks density for the entire Lobe-2, we generated a truncated construct deleting Lobe-2 (residues 1-460). This fragment, containing only the CC domain and Lobe-1, was sufficient to recapitulate the observed dimer-hexamer equilibrium (Supplementary Fig. S6C-F). Furthermore, although ATPgS was included during complex preparation, no density corresponding to bound nucleotide was observed, consistent with HELLS adopting an inactive conformation in the hexameric state.

### ATPase Lobe 2 of HELLS is conformationally flexible and dynamic

To investigate the missing Lobe-2, we optimized the cryo-EM processing strategy by re-extracting particles using a larger box size and reconstructing the map without imposing symmetry (Supplementary Fig. S2). At high contour thresholds, the resulting map closely resembled the D3symmetric reconstruction (Fig. 5A, left panel). However, at lower contour thresholds, substantial density appeared (R1, R2, R3 and R4 in Fig. 5A, right panel), likely corresponding to the previously unresolved Lobe-2 and the CC2-Lobe-1 linker region. DeepEMhancer-sharpened map (50) of the same reconstruction showed improved quality of densities (Fig. 5B). The R1 density was sufficiently well-defined to allow docking of complete AlphaFold-predicted model of Lobe-2 (52) of HELLS into one protomer (Fig. 5C).

**Figure 5.**
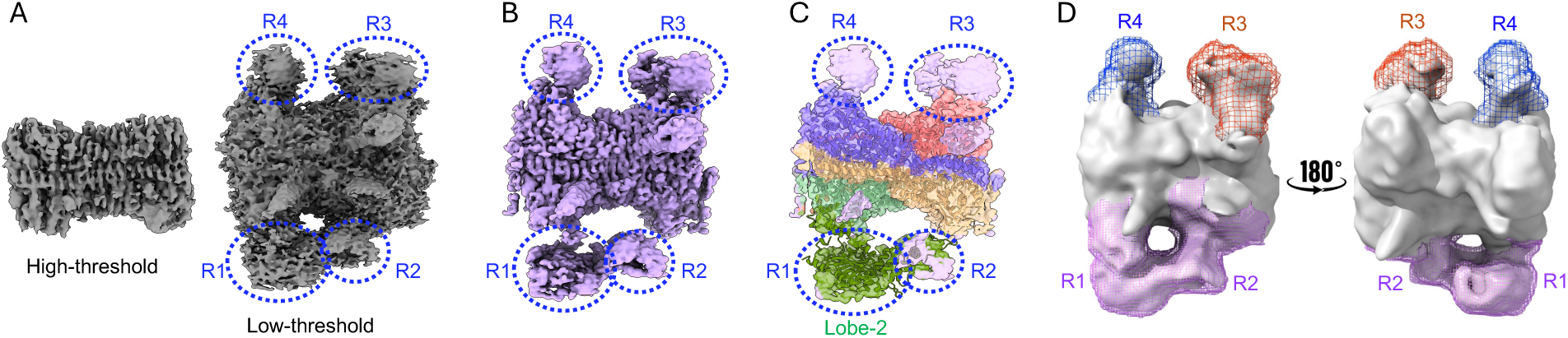
Conformational flexibility and variability of HELLS Lobe-2. **(A)** Representative cryo-EM map of C1 reconstruction shown at high threshold (left panel) and a lower threshold (right panel), revealing additional unmodeled density corresponding to regions (R1, R2, R3 and R4). **(B)** DeepEMhancer-processed map of the same reconstruction, showing improved quality of densities. (**C**) Docking of the AlphaFold-predicted Lobe-2 model into the R1 density region. (**D**) Orthogonal views of the focused masks used for localized refinement of regions R1-R2 (magenta), R2 (orange), and R3 (blue) (Supplementary Fig. S3 and S4).

To further characterize these flexible regions, we performed 3D classification without alignment and conducted 3D variability analysis, each using a focused mask around R1-R2, R3 or R4 (Fig. 5D, Supplementary Fig. S3, Supplementary Movie S1), followed by non-uniform refinement of each class in cryoSPARC (49). These analyses revealed substantial domain motions of Lobe-2 (R1) relative to Lobe-1, pivoting around the flexible linker connecting the two lobes (Supplementary Fig. S4 and Movie S2). In addition, R3-R4 displayed continuous and correlated motions, transitioning between conformations in which these regions are spatially separated and conformations in which they move close together (Supplementary Fig. S3 and Movie S1). Together, these results indicate that the heterogeneity observed in the reconstruction is intrinsic to Lobe-2 in the hexameric assembly.

### Structural comparison with DDM1

We extracted a single HELLS protomer from the hexamer assembly and superimposed it onto the recently reported structure of the DDM1-nucleosome complex (59), which lacks the corresponding N-terminal CC domain (59) (Supplementary Fig. S7A). Lobe-1 from the two structures aligned well, with a root-mean-squared-deviation of 1.4 Å and 54% sequence identity across residues 208-457 (Supplementary Fig. S8). In contrast, Lobe-2 adopts markedly different conformations in the two proteins. In DDM1, Lobe-2 forms a closed arrangement with Lobe-1 that clamps onto nucleosomal DNA. In HELLS, however, Lobe-2 (docked using AlphaFold-fold predicted model) remains in a dynamic and open configuration within the hexamer, suggesting that DNA engagement would require Lobe-2 to rotate toward and closed onto Lobe-1.

The superimposition also showed that the N-terminal CC domain of HELLS is positioned adjacent to nucleosomal DNA. A key distinction between HELLS and DDM1 lies in the sequence composition of their N-terminal CC domains (Supplementary Fig. S7B). Across residues of 24-100, human HELLS and mouse Lsh share 95% sequence identity (only 4 of 77 residues differ), whereas DDM1 shares only 9 identical residues (12% identify) with HELLS in this region (Supplementary Fig. S7B). The charged and hydrophobic residues that mediate HELLS intra- and inter-molecular interactions are not conserved in DDM1 and, in several positions, are replaced with residues of opposite charges. Therefore, the autoinhibitory mechanism identified for HELLS is unlikely to apply to DDM1.

### HELLS-CDCA7-DNA complex

It has been proposed that CDCA7 relieves HELLS autoinhibition (39), as CDCA7 contains a predicted long a-helix (residues 112-130) that may compete with Lobe-1 for binding to the same CC2 surface. To test this model, we generated an N-terminal HisMBP-CDCA7 fusion bearing a C-terminal 6xMyc tag. Expression in baculovirus-infected Sf9 insect cells greatly improved protein yield and solubility compared with *E. coli*, and the C-terminal Myc tag enabled purification of full-length CDCA7, which is otherwise truncated when expressed in bacteria (41). When mixed with HELLS (in either dimeric or hexameric form), CDCA7 co-eluted with HELLS during size-exclusion chromatography, indicating stable complex formation (Fig. 6A).

**Figure 6.**
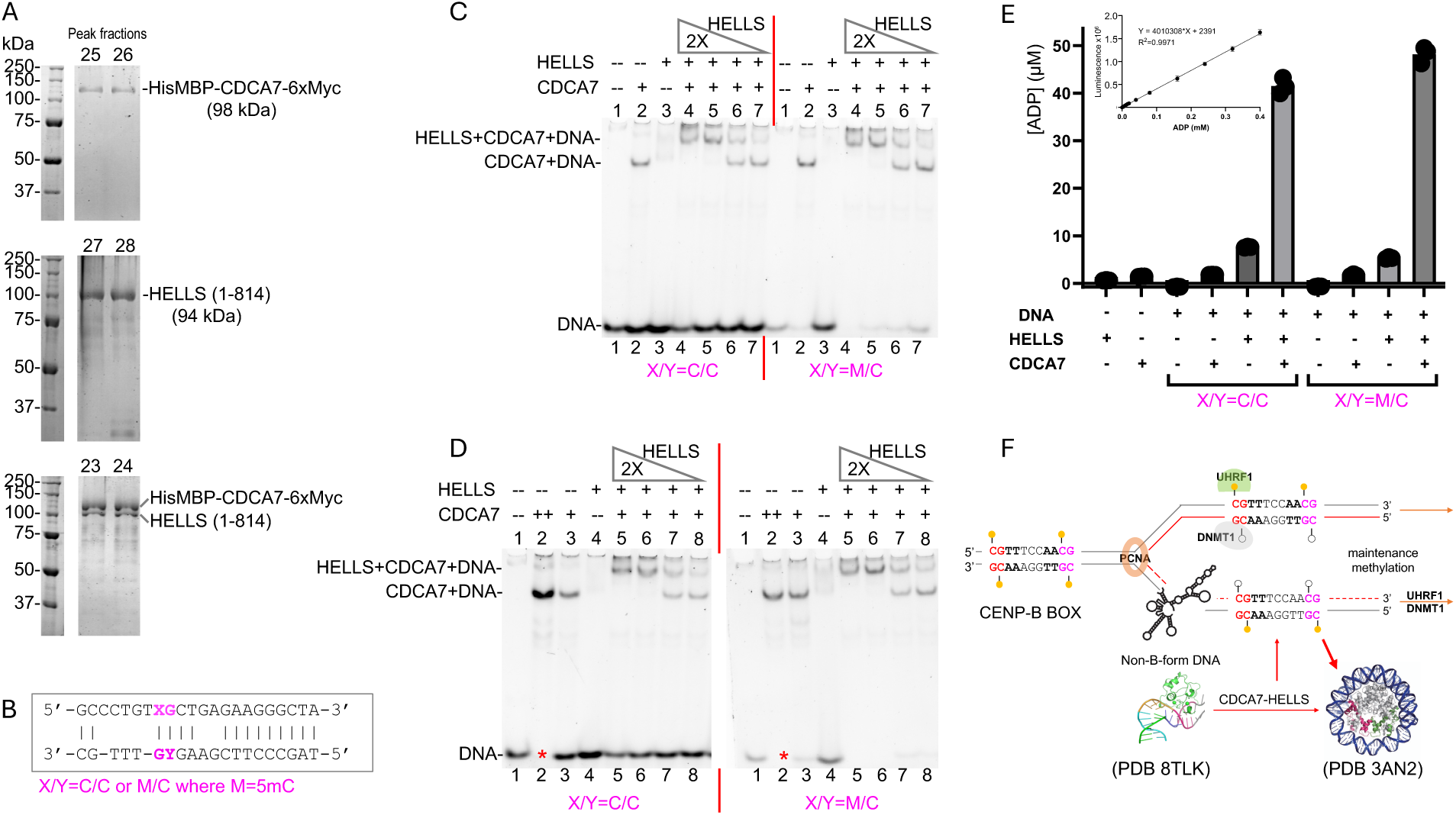
HELLS-CDCA7 interactions in the presence of a short non-B-form DNA substrate. (A) Protein samples used in this study, including peak fractions (labeled) eluted from a Superdex 200 10/300 GL column of HisMBP-CDCA7-6xMyc fusion, HELLS (1-814), and the mixture of both proteins. (B) Non-B-form DNA substrate containing a CpG site (X/Y=C/C). Methylation of the top strand generates a hemi-methylated CpG (X/Y=M/C, M is 5-methylcytosine or 5mC). (C) EMSA using the non-B DNA probe (50 nM, lane 1), CDCA7 (125 nM, lanes 2 and 4-7), HELLS (500 nM, lanes 3-4; twofold serial dilutions in lanes 5-7). (D) EMSA under similar conditions with 50 nM DNA (lane 1), CDCA7 (250 nM, lane 2; 125 nM, lanes 3 and 5-8), and HELLS (500 nM, lanes 4-5; twofold serial dilutions in lanes 6-8). (E) ATPase activity measured with 2.5 μM DNA and 0.25 μM protein. The Y-axis is the ADP concentration produced during the reaction and the inserted is the calibration curve of ADP concentration. (F) Proposed model in which the HELLSCDCA7 complex acts on non-B-form DNA during replication-dependent slippage. During replication-dependent slippage in repetitive DNA, non-B structures can form, leading to misaligned or self-annealed repeats and replication pausing. Under normal conditions without slippage, DNMT1/UHRF1 is sufficient to maintain DNA methylation. When non-B structures arise, the HELLS-CDCA7 complex recognizes hemi-methylated CpG – a replication-specific feature – binds these structures and the helicase activity of HELLS unwinds and converts them back to canonical B-DNA. This restored duplex DNA can then be methylated by DNMT1/UHRF1, reestablishing nucleosome-bound heterochromatin.

To assess DNA binding, we used a short, specific non-B DNA substrate containing a CpG dinucleotide, and its hemi-methylation on one strand, that we recently identified as a CDCA7binding element (41) (Fig. 6B). Electrophoretic mobility shift assay (EMSA) showed that the CDCA7 fusion readily binds DNA, as expected (lane 2 in Fig. 6C). However, HELLS alone did not produce a detectable shift (lane 3 in Fig. 6C), despite the strongly basic surface features of the HELLS hexamer (Fig. 4C). In contrast, the CDCA7-HELLS mixture produced a slower-migrating complex (lanes 4-7), consistent with formation of a ternary HELLS-CDCA7-DNA assembly.

Formation of the super-shifted complex was dependent on the HELLS:CDCA7 ratio. At sub-stoichiometric HELLS (HELLS:CDCA7 £ 1:1), two distinct bands appeared – one corresponding to CDCA7-DNA and the other to the HELLS-CDCA7-DNA complex (lanes 6 and 7). Increasing HELLS beyond equimolar ratios eliminated the CDCA7-DNA band and enhanced the HELLS-CDCA7-DNA complex (lanes 4 and 5). Similar trends were observed using hemimethylated CpG DNA (right portion of Fig. 6C). In a replicate experiment, a two-fold increase in CDCA7 resulted in complete DNA shifting (lane 2 in Fig. 6D). Notably, a previous study in Xenopus egg extracts showed that HELLS and CDCA7 form a stoichiometric complex on chromatin (42).

We next measured ATPase activity following an established protocol (39). Neither HELLS nor CDCA7 alone exhibited ATPase activity. The N-terminal HisMBP tag on CDCA7 had negligible effects on ATPase measurements using the same assay (39). HELLS showed weak but measurable activity in the presence of DNA, consistent with previous reports showing that mouse Lsh has detectable, albeit low, DNA-dependent ATPase activity (11). In contrast, the HELLSCDCA7-DNA ternary complex displayed a ∼5-fold increase in ATPase activity (Fig. 6E). As reported previously (39,42), HELLS ATPase activity is significantly enhanced only when both DNA and CDCA7 are present. A unique aspect of our study is that a non-B-form DNA containing a CpG dinucleotide – rather than canonical Watson-Crick duplex DNA used in prior studies – serves as the stimulatory substrate for HELLS ATPase activation.

## DISCUSSION

Our study provides a detailed structural and biochemical characterization of HELLS, uncovering a previously unrecognized autoinhibitory mechanism and oligomeric architecture that regulates its activity. We show that HELLS assembles into trimers of dimers through two distinct interfaces formed by the CC1 and CC2 helices. Binding of CDCA7 – presumably via its predicted a-helix – disrupts this auto-assembly of HELLS and promotes formation of the HELLS-CDCA7-DNA ternary complex, which exhibits the highest ATPase activity under the conditions tested.

Addition of non-B-form DNA leads to partial disassembly of the hexamer, as evidenced by the measurable DNA-stimulated ATPase activity of HELLS, suggesting that DNA with specific topology may weaken otherwise inert protomer-protomer contacts. This mode of regulation is biologically relevant, as HELLS and CDCA7 are enriched at centromeric and heterochromatic regions, where repetitive DNA sequences frequently adopt non-B-form conformations (60–64). CDCA7 likely acts as an adaptor that links the non-B-form DNA recognition (41) to the CC-domain interface, thereby releasing HELLS from its autoinhibitory state and stabilizing the rearranged HELLS-CDCA7-DNA assembly.

Importantly, the CGTTT sequence (Fig. 6B) required for CDCA7 recognition of non-Bform DNA (41) is embedded within the CENP-B box – a well-characterized DNA motif present in the centromeric repeat units (65). This supports a model in which the HELLS-CDCA7 complex cooperatively targets specialized DNA structures and sequences within repetitive chromatin regions. Because the CDCA7-binding motif contains a CpG dinucleotide – subject to DNA methylation – and CDCA7 preferentially binds hemi-methylated CpG (39–41), this mechanism provides a direct link to the centromeric DNA hypomethylation observed in ICF patients carrying HELLS and CDCA7 mutations (21).

Beyond its nucleosome-sliding activity on canonical duplex DNA (which require neither CpG sequence nor methylation nor participation of DNA repair factors), it has not escaped our attention that the HELLS-CDCA7 complex also resolves (unwinds) non-B-form DNA structures, particularly those arising during replication-dependent slippage that causes repeated sequences to misalign (66). By converting these non-B structures back to duplex DNA, the HELLS-CDCA7 complex may facilitate access by the DNMT1/UHRF1 maintenance machinery, thereby restoring DNA methylation and heterochromatin integrity (Fig. 6F). This restoration and repair activity appears tightly regulated - ATP-dependent, temporally restricted to S-phase, and spatially concentrated at centromeres. Consistent with this model, HELLS depletion in human U2OS cells leads to degradation of nascent DNA at stalled replication forks and increased genomic instability (67). Moreover, in HEK293 cells, CDCA7 and HELLS coimmunoprecipitate with the classical nonhomologous end-joining proteins Ku80 and Ku70 (43), supporting a coordinated role in maintaining genome stability and DNA methylation, potentially through resolving non-B form DNA intermediates. We note that ATRX, another chromatin-remodeling protein, protects against replication stress in part by resolving G-quadruplex DNA structure (68) or other DNA structures related to telomeric repeats (69,70). Similarly, two RecQ-family helicases WRN (Werner syndrome helicase) and BLM (Bloom syndrome helicase) function as non-B DNA-structure-specific helicases (71–74), underscoring a broader paradigm in which specialized genome-maintenance factors protect stability by resolving alternative DNA conformations.

In summary, this study defines the structural basis of HELLS autoinhibition and its potential activation, revealing a regulatory mechanism in which coiled-coil–mediated oligomerization imposes Lobe-1 rigidity and Lobe-2 flexibility. By coupling its oligomeric state to ATPase control, the HELLS-CDCA7 complex employs a distinct allosteric strategy that targets non-B-form DNA – often associated with DNA damage – setting it apart from other SNF2 remodelers. These insights provide a foundation for future work on how mutations in HELLS and CDCA7 disrupt this regulatory balance in disorders such as ICF syndrome and cancer.

## DATA AVAILABILITY

The cryo-EM density maps and their respective atomic coordinate files have been deposited to the Electron Microscopy Data Bank (EMDB) and Protein Data Bank (PDB) under the following accession codes: D3 symmetry map and structure model (EMD-73693, PDB 9Z04), D3 with symmetry expansion (EMD-73694, PDB 9Z05), C1 symmetry map and structure model (EMD73695, PDB 9Z06), C1 symmetry map with additional (R1-R4) densities (EMD-73696), and R1focused map (EMD-73697). The raw data had been deposited to Electron Microscopy Public Image Archive (EMPAIR-13146).

## ACKNOWLEDGEMENTS

We thank Nalahari Akkladevi and Paul Leonard (MD Anderson Cancer Center) for access to the Vitrobot, the cryo-EM facility at Baylor College of Medicine for microscope access during grid optimization, and the Recombinant Protein Production and Characterization Core (Baylor) for analytical ultracentrifugation. We thank Venkata Mallampalli (UTHealth Cryo-EM Core) for Krios data collection. High-resolution data were collected with support from Kasahun Neselu at the National Center for CryoEM Access and Training (NCCAT) and the Simons Electron Microscopy Center (Project ID: NCCAT-BAG-GK210101). We acknowledge ANVIL-ACCESS/Rosen Center for Advanced Computing (Project ID: BIO240101), Nalahari Akkladevi, and the Seadragon highperformance computing system at MDACC for CryoSPARC and RELION computational resources. We thank Swanand Hardikar (MDACC) for technical assistance, and Akeo Shinkai and Yoichi Shinkai of RIKEN (Japan) for discussion.

## AUTHOR CONTRIBUTIONS

G.K. performed cryo-EM and biophysical experiments, purified HELLS proteins, and contributed to writing and editing of the manuscript. R.R. generated HELLS constructs for *E. coli* expression, purified HELLS proteins, and performed DLS. J.L. produced the CDCA7 construct in Sf9 cells, purified CDCA7 and carried out EMSA and ATPase activity assays. J.R.H. assisted with structural refinement and purified HELLS (1–460) truncation proteins. X.Z. contributed to discussions, supervision and project administration. Y.G. supervised and guided G.K. in cryo-EM data processing. T.C. initiated the study, provided Lsh constructs, and assisted with manuscript editing. X.C. designed and organized the study, contributed to manuscript writing and editing, conceptualization, and funding acquisition.

