## supplementary for "Structure of human lymphoid-specific helicase HELLS in its autoinhibitory state"

Figures S1-S8

Table S1

Movies S1-S2

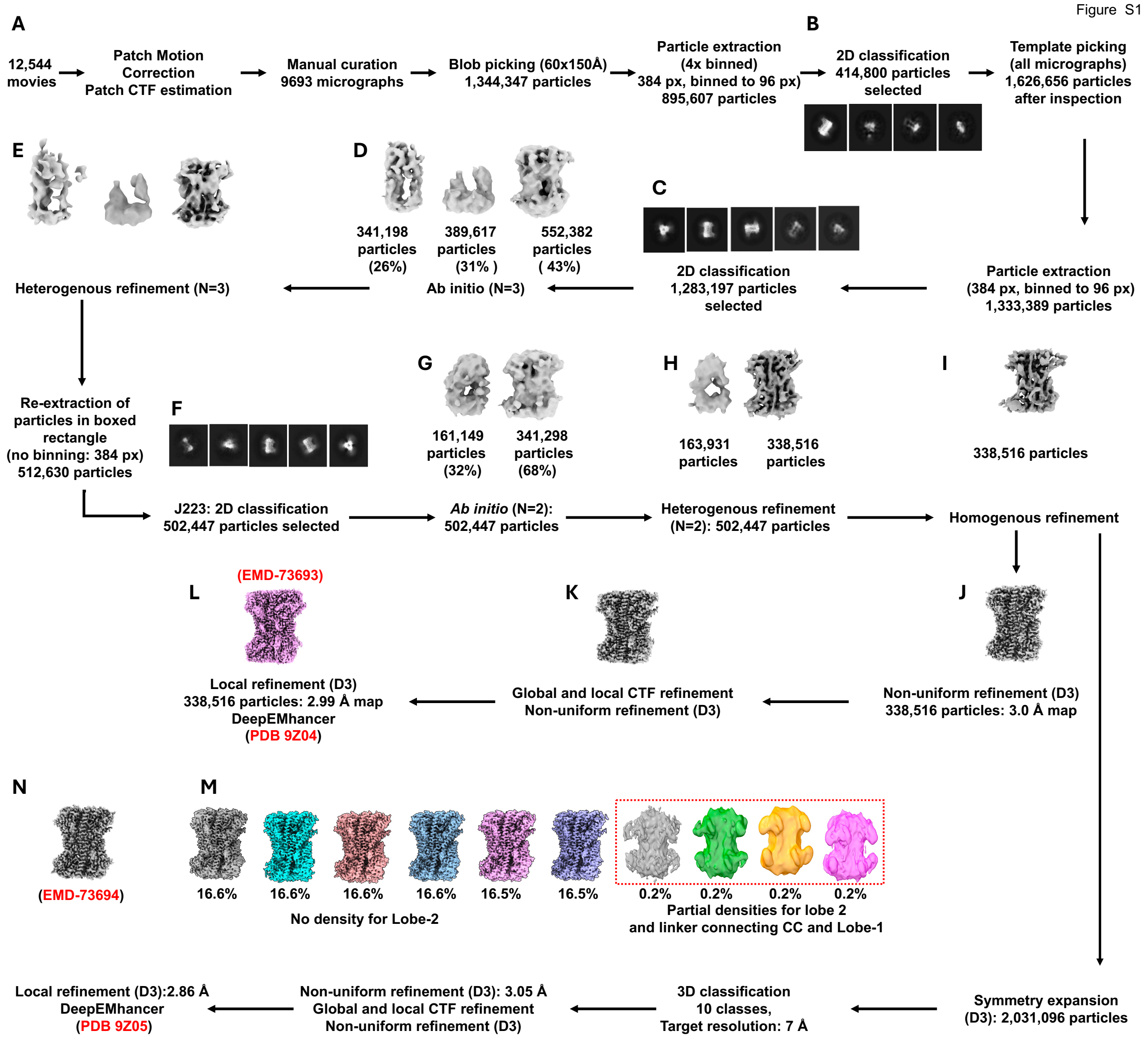


**Supplementary Figure S1. Workflow of D3 reconstruction.**

Micrographs were imported into cryoSPRAC (v4.5.3) (1) (Fig. S1A) and were motion-corrected, dose-weighted, and CTF (Contrast Transfer Function)-estimated. After manual curation, 9693 (out of 12,544 or 77%) micrographs were retained for further analysis for CTF fit resolution, ice thickness, astigmatism, and defocus. The micrographs were split into four sets, one subset was used for blob-based particle picking [a minimal diameter 60 Å and maximum diameter of 150 Å, elliptical shape, and a minimum separation distance of 2 Å diameter], 2D classification [a box size of 384 pixels x 384 pixels (4x binned)] (Fig. S1B), and generation of template, which were than applied to the remaining three subsets. In total, 1,333,389 particles were extracted [a 384-pixel box size (4x binned)] for 2D classification (Fig. S1C), *ab initio* 3D reconstruction (n=3) (Fig. S1D), and heterogenous refinement (Fig. S1E), yielding a best class of 552,382 particles. These were re-extracted (2x binned), further 2D-classified (502,447 particles) (Fig. S1F) into two classes of ab initio models (Fig. S1G), further refined through heterogeneous rounds (Fig. S1H), resulting in a well-resolved class of 338,516 particles. Homogenous (Fig. S1I) and non-uniform refinement (Fig. S1J) using D3 symmetry, followed by global and local CTF refinement (Fig. S1K), local refinement with D3 symmetry (Fig. S1L), followed by 'Fit Spherical Aberration', Fit Tetrafoil', and 'Fit Anisotropic Magnification', produced a final map of 2.99 Å resolution [FSC=0.143 criterion (2)] (PDB 9Z04). The reconstruction was post-processed using DeepEMhancer sharpening (3) at COSMIC2 (4).

To resolve asymmetric features within the D3-symmetric HELLS complex, 338,516 particles from homogenous refinement (Fig. S1I) were symmetry-expanded and subjected to 10-class 3D classification and non-uniform-refinement (Fig. S1M), followed by global and local CTF refinement. Local refinement with 'Fit Spherical Aberration', 'Fit Tetrafoil', and 'Fit Anisotropic Magnification' yielded a map of 2.86 Å [FSC=0.143 criterion (2)] (PDB 9Z05). The final reconstruction was sharpened with DeepEMhancer (3) and processed on the COSMIC2 platform (4) (Fig. S1N).

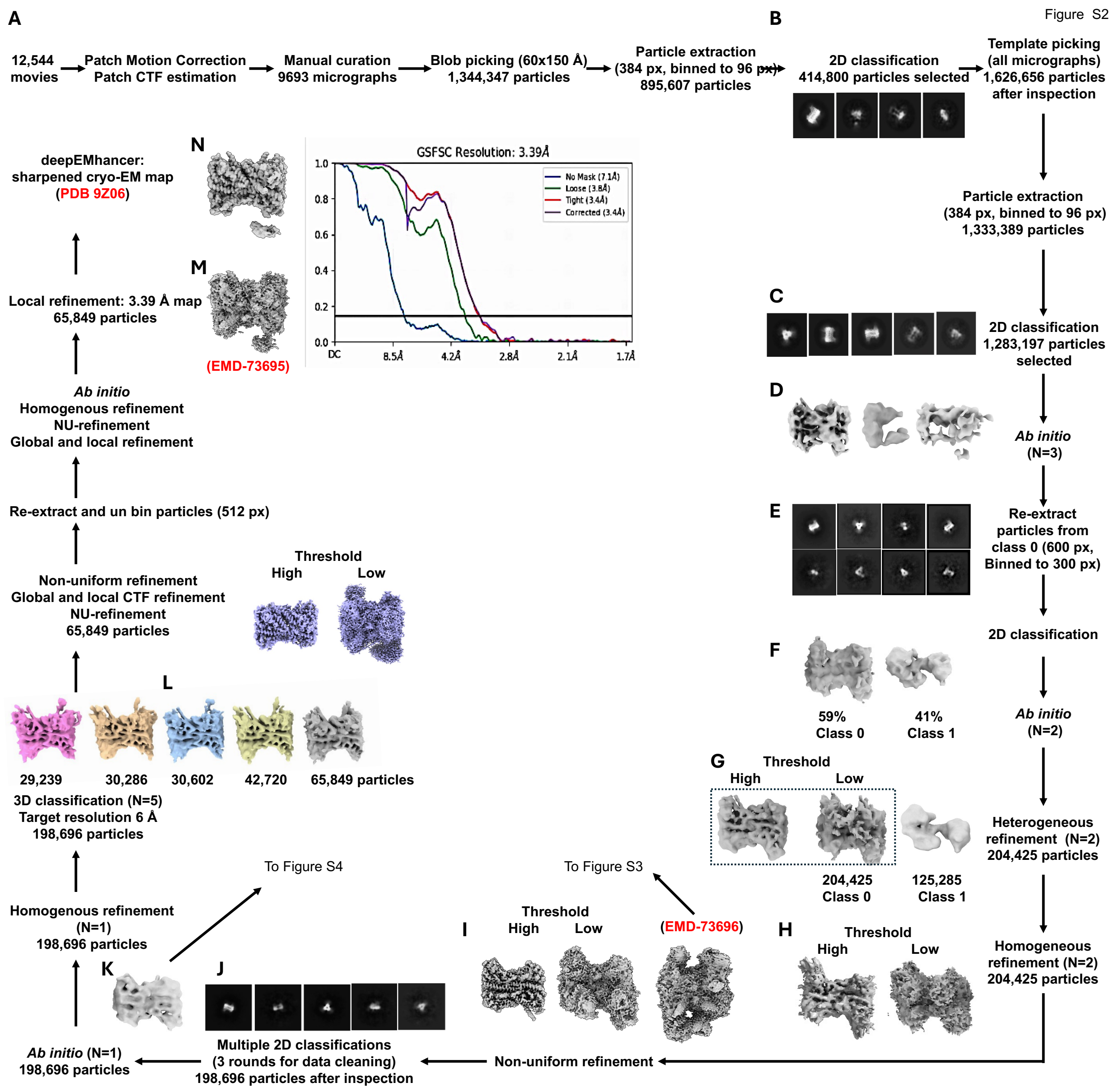


**Supplementary Figure S2. Workflow of C1 reconstruction.**

The initial processing steps (Fig. S2A-D) followed those described in Supplementary Fig. S1. Particles from the best *ab initio* class were re-extracted using a larger 600x600 pixel box (2x binned) (Fig. S2E), subjected to multiple rounds of 2D classification, and used to generate two *ab initio* models (Fig. S2F) and heterogenous refinement (Fig. S2G). The best class underwent homogenous refinement (Fig. S2H) and non-uniform refinement in C1 symmetry (Fig. S2I).

Additional 2D classification removed suboptimal particles (Fig. S2J), yielding a curated particle set for a single ab initio model and homogenous refinement (Fig. S2K). Due to anisotropy, 3D classification (n = 5 classes, target filter 6 Å) (Fig. S2L) and non-uniform refinement were performed, and the best class was re-extracted unbinned for further processing. Global and local CTF refinement followed by non-uniform and local refinement in C1 symmetry produced the map in cryoSPARC (Fig. S2M), with 'Fit Spherical Aberration', 'Fit Tetrafoil', and 'Fit Anisotropic Magnification' options enabled. The final map was subsequently sharpened using DeepEMhancer (3) on the COSMIC2 platform (4) (Fig. S2N).

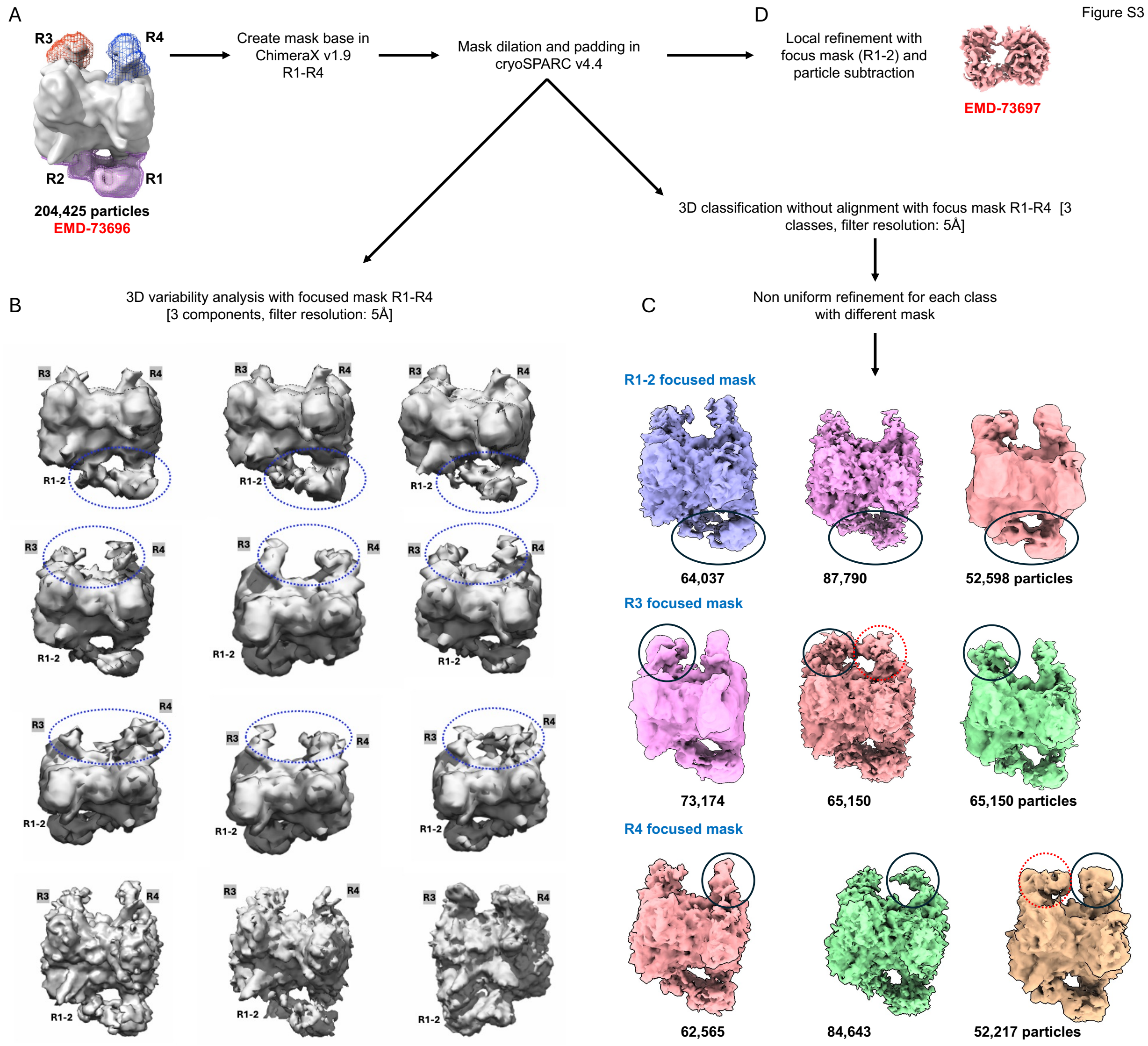


**Supplementary Figure S3**. **Workflow for focused refinement and analysis of flexible regions (R1-R4).** To assess flexibility in the EMD-73696 reconstruction (Fig. S3A), regions R1–R2, R3, and R4 were segmented in ChimeraX v1.9 to create initial masks, which were then dilated and padded in cryoSPARC v4.5.3. Using these focused masks, 3D variability analysis (3DVA) captured continuous motions of each flexible lobe (Fig. S3B). In parallel, 3D classification without alignment followed by non-uniform refinement identified discrete conformational states for R1-R2, R3, and R4 (Fig. S3C). Focused refinement of R1–R2 yielded a sharpened map (EMD-73697) (Fig. S3D).

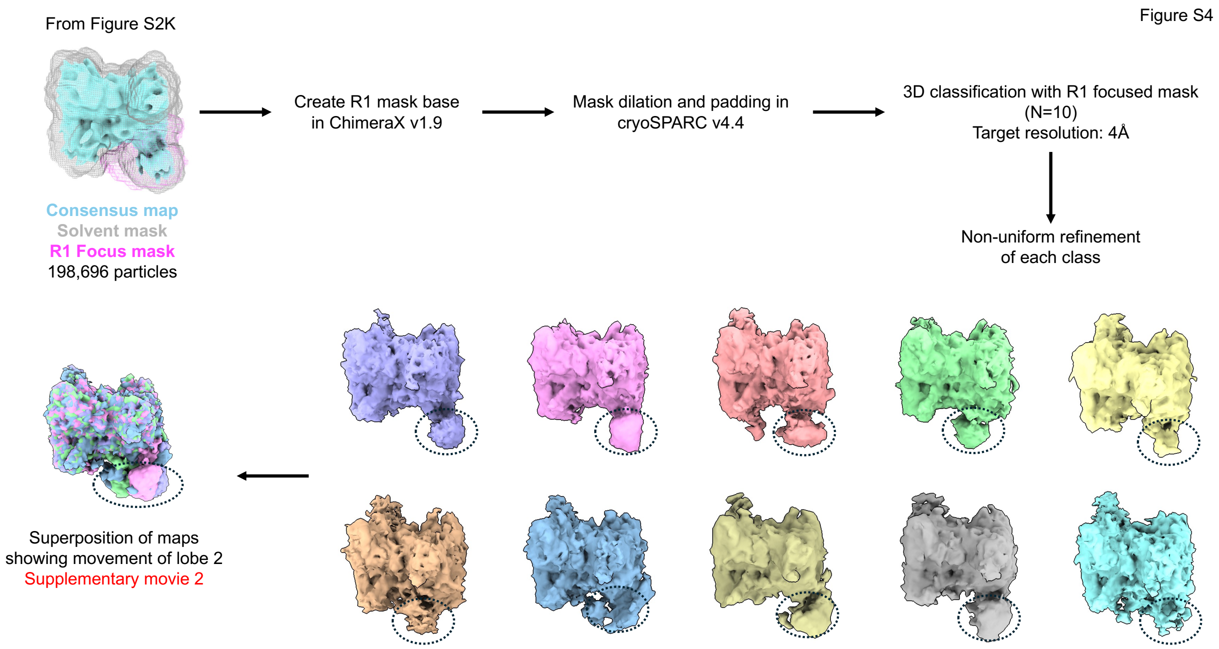


**Supplementary Figure S4**. **Focused refinement of R1 region and analysis of flexible Lobe-2.** A focused mask for R1 was generated in UCSF ChimeraX (5) and used for 3D classification into 10 classes in cryoSPARC (1), with a target filter resolution of 4 Å, followed by non-uniform refinement of each class.

**Supplementary Movie 1.** Three-component 3D variability analysis (3DVA) of the EMD-73696 particle set. Movie 1A shows continuous conformational motions within flexible regions R1–R2, while Movie 1B captures coordinated and independent movements of regions R3 and R4.

**Supplementary Movie 2.** Focused 3D classification of the R1 region reveals its dynamic and heterogeneous conformational behavior.

Figure S5

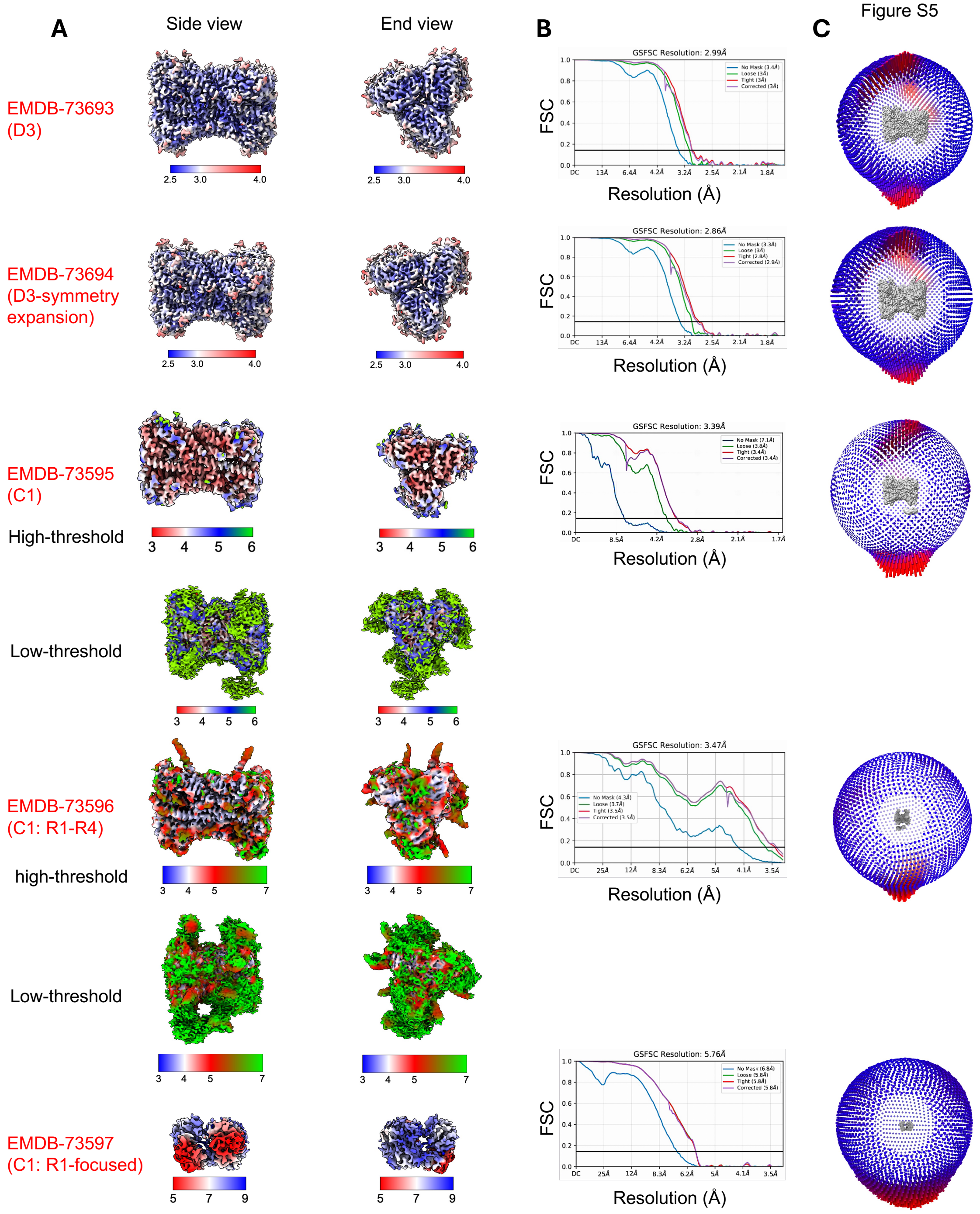


**Supplementary Figure S5. Local resolutions and angular distribution analyses for cryo-EM reconstruction**. (**A**) Orthogonal side and end views of the five cryo-EM maps, with corresponding EMDB accession numbers. (**B**) Fourier shell correlation (FSC) curves for each reconstruction. (**C**) 3D angular distribution heatmaps for each reconstruction.

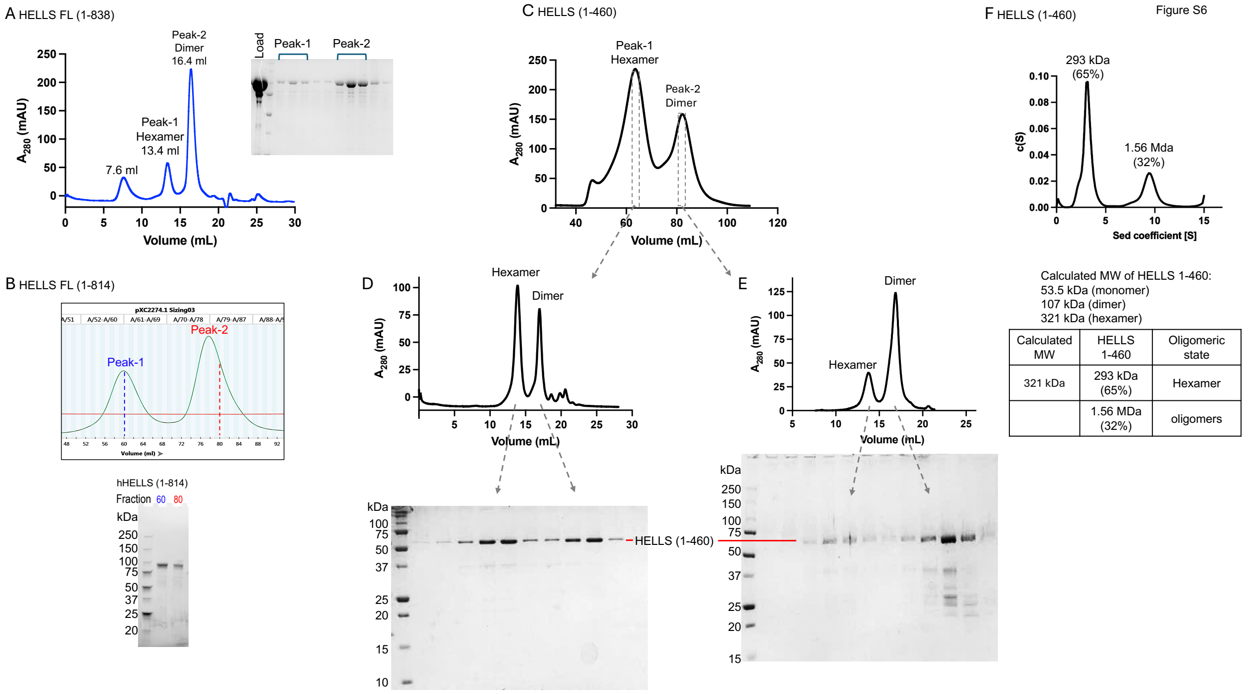


**Supplementary Figure S6** (related to Figure 1). **Purification of recombinant HELLS constructs of three different lengths.** (**A**) SEC profile of full-length HELLS (1-838) on a Superose6 increase 10/300GL column. (**B**) SEC profile of HELLS (1-814) on Superdex S200 16/60 column (Cytivia). SDS-PAGE analysis of fractions 60 (peak-1) and 80 (peak-2) is shown below. These two fractions were used for cryo-EM sample preparation. (**C-E**) SEC profile of HELLS (1-460) on Superdex S200 16/60 column (Cytivia). Peak fractions from panel C were re-run on Superose6 increase 10/300 GL in panels D and E. Corresponding SDS-PAGE gels are shown below each profile. (**F**) SV-AUC sedimentation profile of HELLS (1-460), with derived molecular masses tabulated below.

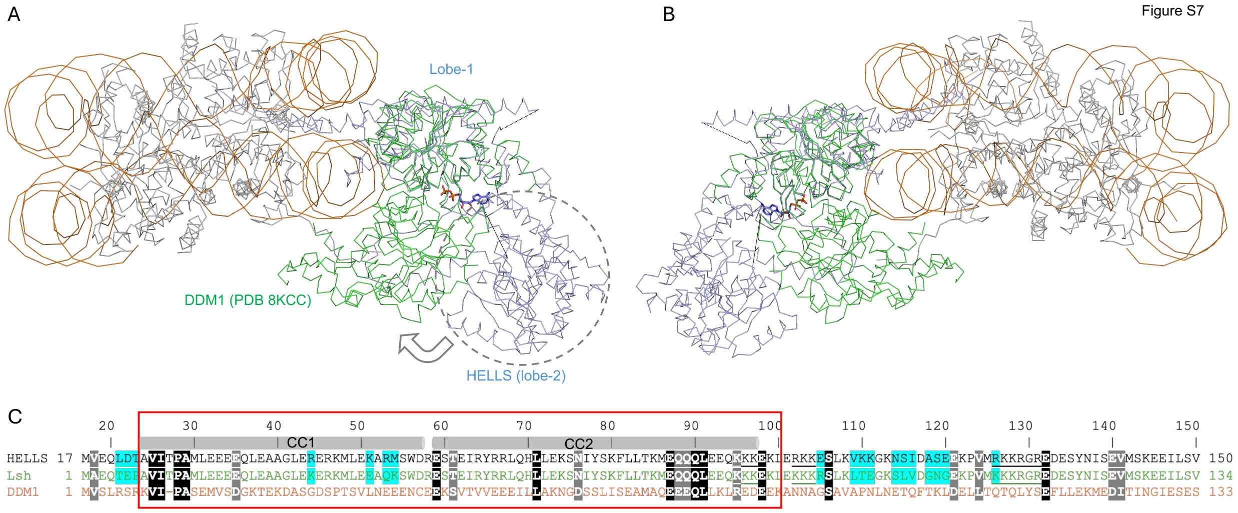


**Supplementary Figure S7. Comparison of human HELLS with the *Arabidopsis thaliana* homolog DDM1.** (**A**-**B**) Two opposite views of the superimposition of HELLS (light blue) onto nucleosome-bound DDM1 (green; PDB 8KCC (6)), aligned on Lobe-1. (**C**) The N-terminal sequence alignment of human HELLS, mouse Lsh, and *Arabidopsis* DDM1 including the CC domain regions. Although not conserved and not visible in the structure, deletion of DDM1 residues 1-132 increases DDM1 ATPase activity, indicating an N-terminal autoinhibitory function (7).

Figure S8

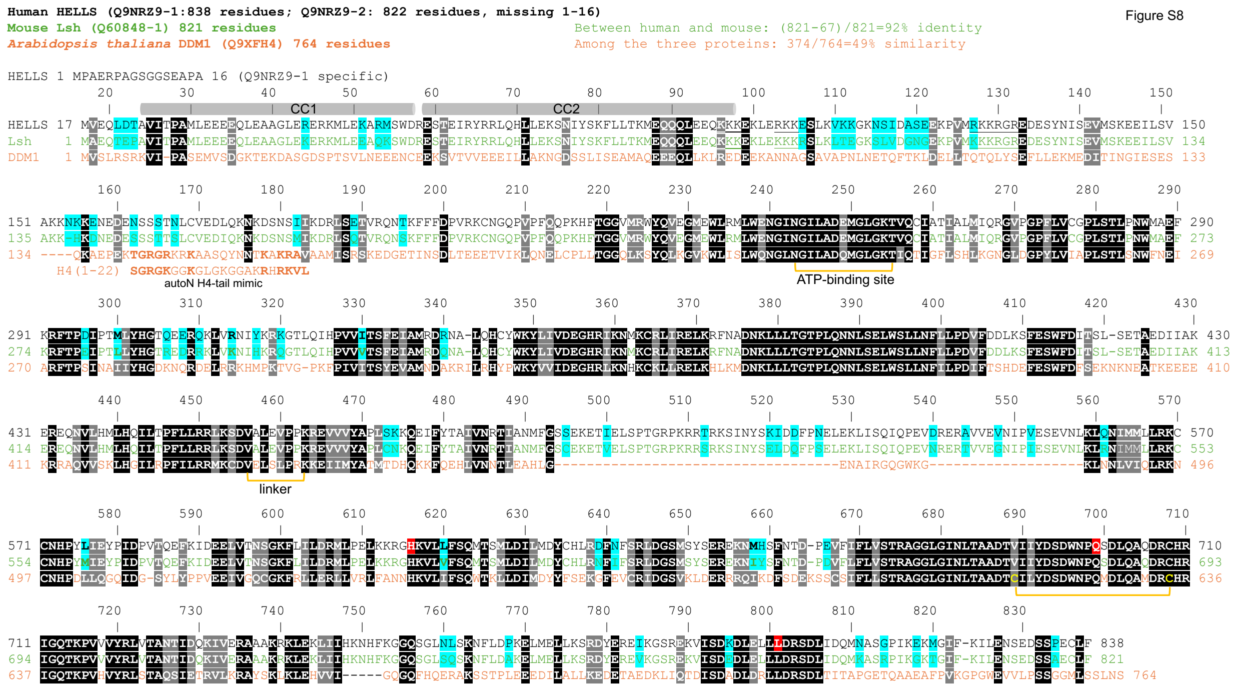


**Supplementary Figure S8**. **Sequence alignment of three helicases** - human HELLS (UniProt Q9NRZ9-1), mouse Lsh (UniProt Q60848-1), and *Arabidopsis* DDM1 (UniProt Q9XFH4). The alignment is arranged to minimize insertions and deletions. Full-length HELLS contains an additional N-terminal 16 residues that are absent in the shorter isoform Q9NRZ9-2. DDM1 is the smallest of the three and contains a three-residue deletion between residues 133-134, a large 70-residue deletion between 484-485, and a five-residue deletion between 673-674. Invariant residues shared by all three enzymes are shown as white letters on a black background, whereas conserved residues are shown as white letters on a light grey background (A, V, L, I, and M; S and T; E and D; Q and N; K and R). The HELLS CC1 and CC2 α-helices are annotated above the alignment. Validated ICF variants in HELLS are highlighted in red - H616R and Q699R (missense) and L801del (deletion). A disulfide bond (S-S) between C615 and C634 of DDM1 (7) is indicated by a bracket. Full-length DDM1 binds with unmodified (deacetylated) H4 peptides (7), which shares sequence similarity with DDM1 residues 141-162.

**Supplementary Table S1**. Cryo-EM data collection, model refinement, and validation statistics

| Construct | HELLS (1-814) | | | | |
|---|---|---|---|---|---|
| Symmetry imposed | D3 | | C1 | | |
| | | Symmetry expansion | | Map with R1-to-R4 densities | R1-Focused map |
| Composition | CC-Lobe1 | CC-Lobe1 | CC-Lobe1 | | |
| EMDB | 73693 | 73694 | 73695 | 73696 | 73697 |
| PDB | 9Z04 | 9Z05 | 9Z06 | - | - |
| Resolution (Å) | 2.99 | 2.86 | 3.39 | 3.47 | 5.76 |
| **Data collection and processing** | | | | | |
| Electron Microscope | Krios | | | | |
| Camera | Gatan K3 | | | | |
| Magnification | 105,000X | | | | |
| Voltage (kV) | 300 | | | | |
| Dose rate (e-/Å$^2$/s) | 29.26 | | | | |
| Total dose (e-/Å$^2$) | 58.51 | | | | |
| Frame rate (ms) | 40 | | | | |
| Total frames | 50 | | | | |
| Defocus range (*µm)* | -0.7 to -2.0 | | | | |
| Pixel size (Å) | 0.4135 | | | | |
| Total micrographs (#) | 12,545 | | | | |
| Software | CryoSPARC v4.5.3 and RELION | | | | |
| Initial particles (#) | 1,626,656 | | | | |
| Particle # in final map | 338,516 | 337,716 | 65,849 | 204,425 | 65,849 |
| FSC threshold | 0.143 | | | | |
| **Refinement** | | | | | |
| Non-hydrogen atoms | 16,158 | 15,960 | 15,560 | | |
| Protein residues | 1,974 | 1,944 | 1,892 | | |
| **B-factors (Å$^2$)** | | | | | |
| Protein | 46.95 | 42.36 | 88.35 | | |
| **R.m.s deviations** | | | | | |
| Bond lengths (Å) | 0.003 | 0.003 | 0.003 | | |
| Bond angles (°) | 0.4 | 0.4 | 0.6 | | |
| **Validation** | | | | | |
| MolProbity score | 1.2 | 1.3 | 1.9 | | |
| Clash score | 4.0 | 6.0 | 9.1 | | |
| **Ramachandran plot** | | | | | |
| Favored (%) | 98.4 | 98.8 | 98.0 | | |
| Allowed (%) | 1.6 | 1.2 | 2.0 | | |
| Outliers (%) | 0 | 0 | 0 | | |
| Rotamer outliers (%) | 1.0 | 1.0 | 3.2 | | |
